# Structural and mechanistic insight into spectral tuning in flavin-binding fluorescent proteins

**DOI:** 10.1101/2021.01.08.425906

**Authors:** Katrin Röllen, Joachim Granzin, Alina Remeeva, Mehdi D. Davari, Thomas Gensch, Vera V. Nazarenko, Kirill Kovalev, Andrey Bogorodskiy, Valentin Borshchevskiy, Stefanie Hemmer, Ulrich Schwaneberg, Valentin Gordeliy, Karl-Erich Jaeger, Renu Batra-Safferling, Ivan Gushchin, Ulrich Krauss

## Abstract

Determining the molecular origin of spectral tuning in photoactive biological systems is instrumental for understanding their function. Spectral-tuning efforts for flavin-binding fluorescent proteins (FbFPs), an emerging class of fluorescent reporters, are limited by their dependency on protein-bound flavins, whose structure and hence electronic properties, cannot be altered by mutation. To address those shortcomings, we here present the photophysical, computational and structural characterization of structurally uncharacterized blue-shifted FbFPs, carrying a previously described lysine substitution within their flavin-binding pocket. X-ray structures reveal displacement of the lysine away from the chromophore and opening up of the structure as cause for the blue shift. Site-saturation mutagenesis and high-throughput screening, yielded a red-shifted variant, in which the lysine side chain of the blue-shifted variant is stabilized in close distance to the flavin by a secondary mutation, mechanistically accounting for the red shift. Thus, a single secondary mutation in a blue-shifted variant is sufficient to generate a red-shifted FbFP. Using spectroscopy, X-ray crystallography and quantum mechanics molecular mechanics calculations, we provide a firm structural and functional understanding of spectral tuning in FbFPs. We also show that the identified blue- and red-shifted variants allow for two-color microscopy based on spectral separation. In summary, the generated blue- and red-shifted variants represent promising new tools that should find application in life sciences.

## Introduction

Flavin-binding fluorescent proteins (FbFP), engineered from pro- and eukaryotic, light oxygen, voltage (LOV) photoreceptors [1, 2], are promising fluorescent reporter proteins (FPs). Their small size (about 10 kDa for monomeric FbFPs such as the iLOV protein derived from *Arabidopsis thaliana* phototropin photoreceptors [1]) as well as their oxygen-independent fluorescence make them ideal reporters to monitor transport [1, 3] and anaerobic biological processes [4-7]. They are also used as donor domain in Förster Resonance Energy Transfer (FRET)-based biosensors, e.g. for molecular oxygen or pH [8, 9]. Proteins from the Green Fluorescent Protein (GFP) family are very popular for similar applications in life science but have clear disadvantages due to their larger size and their need of sufficient molecular oxygen for chromophore maturation, which often causes failure when used in organisms living under anaerobic conditions. Moreover, FbFPs [1, 2], LOV photosensory domains and other flavin-binding photoreceptors (summarized in [10, 11]) have been utilized as biosensors [8, 9, 12], photosensitizers [13-15] for e.g. correlative light electron microscopy (CLEM) [15], chromophore-assisted light inactivation (CALI) [16] and/or as optogenetic tools for the reversible control of biological processes (recently reviewed in [10]).

Applications, like multi-color imaging, the design of solely FbFP-based FRET biosensors, multi-color CLEM, CALI and LOV-based optogenetics would require the development of spectrally shifted FbFPs or LOV domains. In contrast to GFP-like FPs, whose absorption and fluorescence maxima can be tuned by exchanging certain chromophore constituting amino acids [17], the absorption and fluorescence of FbFPs and other flavoproteins depends on non-covalently bound flavins, whose structure cannot be influenced by mutation. This is a severe drawback of FbFPs, which has so far limited all color-tuning efforts for this important new family of fluorescent reporters.

At present, only blue-shifted FbFPs are available, derived by substituting a fully conserved glutamine (Q489 in iLOV) against valine [18] or lysine [19]. While molecular dynamic (MD) simulations and quantum mechanics molecular mechanics (QM/MM) calculations suggested that e.g. the introduced lysine (K489 in iLOV-Q489K) flips away from the chromophore, accounting for the observed blue shift [19], this theoretical hypothesis remains structurally unproven. Likewise, theoretical predictions suggested that the retention of a charged amino acid, such as lysine, close to the FMN ring system should result in red-shifted spectral properties [19-21], which so far could not be achieved experimentally. In addition, the transferability of the proposed mutations and mechanisms for spectral tuning have so far not been validated for FbFPs, other than iLOV, which however is a prerequisite for widespread implementation.

## Results and Discussion

To verify the transferability of the blue-shifting Q489K mutation, identified for the plant FbFP iLOV [19], we introduced the corresponding Q148K mutation into the recently presented thermostable, CagFbFP protein from thermophilic phototrophic *Chloroflexus aggregans* [22]. In contrast to the monomeric, plant derived iLOV protein, the bacterial CagFbFP protein is a dimer in solution [22]. In terms of sequence, iLOV and CagFbFP share only about 40 % identical amino acid positions, highlighting the distinct origin of CagFbFP. With regard to its flavin-binding site and the overall LOV domain structure, iLOV and CagFbFP are, however, very similar (C-α atom RMSD 0.575 Å over 104 aligned residues) [22]. To study the spectral properties of the CagFbFP-Q148K variant, we heterologously produced the variant in *E. coli* and purified it to homogeneity. The absorption and fluorescence emission spectra of CagFbFP-Q148K are shown in Figure 1, A, B. For comparison, the corresponding spectra of iLOV-Q489K are depicted in Figure 1, C, D.

**Figure 1:**
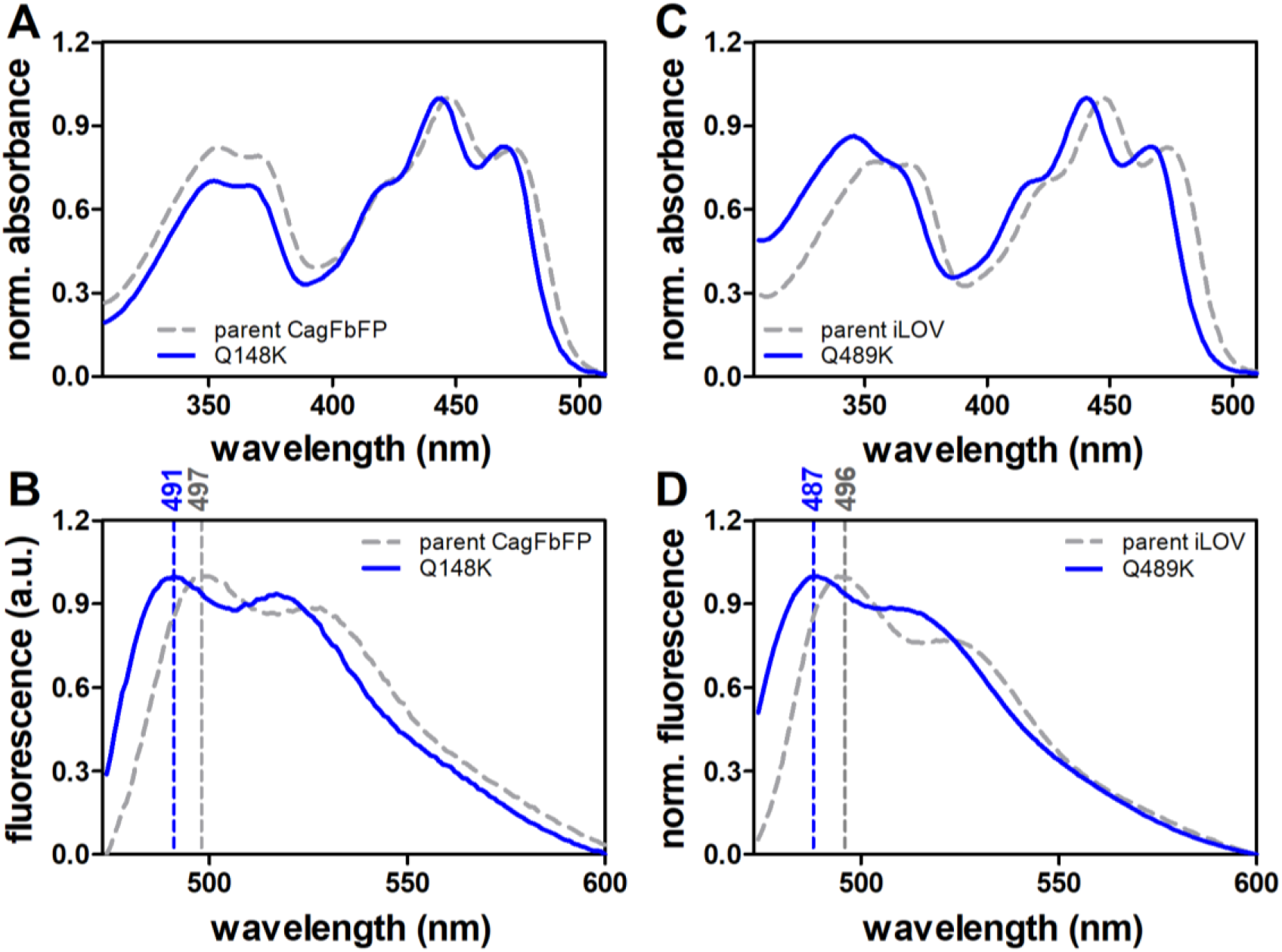
Spectral properties of blue-shifted FbFPs. Absorption (A, C) and fluorescence emission spectra (B, D) of parental CagFbFP and iLOV (grey, dashed line) and their blue-shifted CagFbFP-Q148K/iLOV-Q489K variants (blue, solid line). Dashed lines of the respective color mark fluorescence emission maxima. All spectra are normalized to the corresponding maximum.

Much like iLOV-Q489K (Figure 1, C, D; Table 1), CagFbFP-Q148K possesses a blue-shifted S_0_→S_1_ absorption band with a maximum at around 444 nm (≈ 3 nm blue-shifted relative to parental CagFbFP) as well as a blue-shifted fluorescence-emission maximum (≈ 6 nm blue shift relative to parental CagFbFP; Figure 1, A, B; compare grey dashed line and solid blue line; Table 1). Thus, in conclusion, the spectral effects previously seen in the plant derived, monomeric iLOV-Q489K protein are well reproduced in the thermostable, dimeric CagFbFP protein of bacterial origin, thus hinting at a conserved structural mechanism.

**Table 1:** Photophysical properties of all iLOV and CagFbFP variants

| Protein | $\lambda_{\text{max-absorption}}$<br>(nm) | $\lambda_{\text{max-emission}}$<br>(nm) | shift<br>(nm)<sup>a</sup> | $\Phi_F$ | $\tau_{av}$ / ns |
| --- | --- | --- | --- | --- | --- |
| iLOV | 447 | 496 | - | $0.33 \pm 0.01$ | $4.47 \pm 0.03$ |
| iLOV-Q489K | 441 | 487 | -9 | $0.35 \pm 0.01$ | $4.03 \pm 0.08$ |
| iLOV-Q489K-V392T | 447 | 502 | +6 | $0.33 \pm 0.02$ | $4.48 \pm 0.03$ |
| CagFbFP | 447 | 497 | - | $0.36 \pm 0.01^b$ | $4.53 \pm 0.03$ |
| CagFbFP-Q148K | 444 | 491 | -6 | $0.24 \pm 0.01$ | $3.28 \pm 0.06$ |
| CagFbFP-Q148K/I52T | 450 | 504 | +7 | $0.27 \pm 0.01$ | $3.82 \pm 0.05$ |
| CagFbFP- I52T | 450 | 497 | 0 | $0.38 \pm 0.06$ | $4.42 \pm 0.07$ |

n.d. not determined, <sup>a</sup>: relative to emission maximum of the corresponding parental protein, <sup>b</sup>: data present in [22].

To elucidate the structural basis for the blue-shifted spectral properties we determined the structure of iLOV-Q489K. Tetragonal crystals (space group P4_3_2_1_2; one molecule per asymmetric unit) grew within six weeks, diffracting up 1.45 Å resolution. Data and refinement statistics are summarized in Supplementary Table S1. Globally, the iLOV-Q489K structure is very similar to the structure of parental iLOV (PDB-ID: 4EES; root mean square deviation RMSD 0.6913 Å C-α atoms over residues 104 aligned residues) showing a typical α+β PAS topology with the secondary structure elements in the order Aβ-Bβ-Cα-Dα-Eα-Fα-Gβ-Hβ-Iβ. Surprisingly, the iLOV-Q489K and parental iLOV structures diverge at the C-terminal end following the introduced lysine (K489 in iLOV-Q489K) (Figure 2, A).

**Figure 2:**
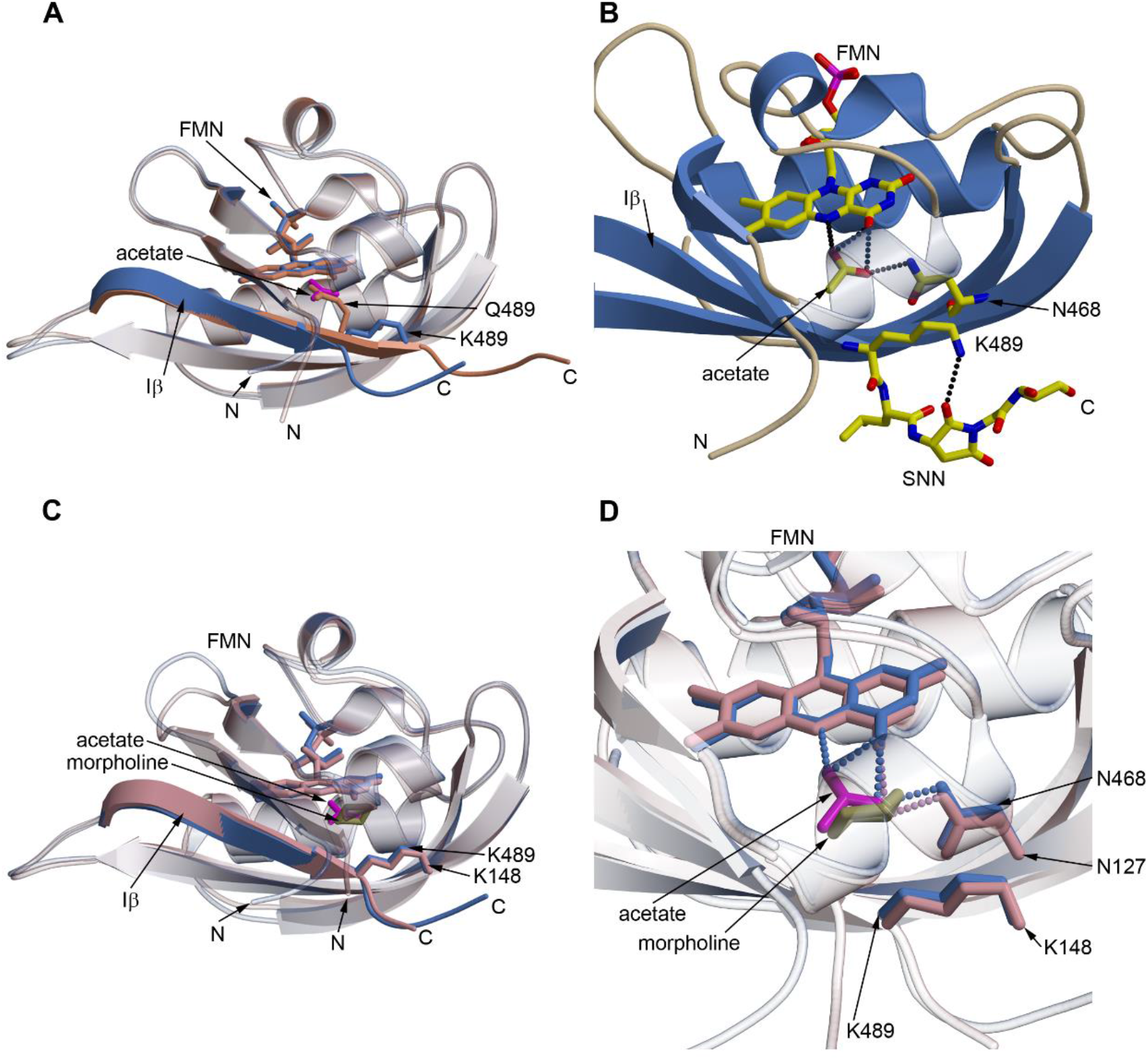
Structural comparison of parental iLOV, iLOV-Q489K and CagFbFP-Q148K. (A) Superposition of parental iLOV (PDB-ID: <u>**4EES**</u>; light salmon) with iLOV-Q489K (cornflower blue). Introduction of Q489K results in a shortening the C-terminal Iβ strand and makes it bend downwards. A buffer molecule fragment [CH_3_COO]^-^ (acetate anion, magenta) occupies the position of the side chain of Q489 in parental iLOV. (B) Structure of iLOV-Q489K. Depicted are the acetate anion with its hydrogen bond network to FMN and N468 (as stick model in standard atomic coloration). The introduced K489 interacts with the modified C-terminus, formed by deamidation of N491 and G492 yielding a succinimide intermediate (SNN L-3-aminosuccinimide). (C) Superposition of iLOV-Q489K (cornflower blue) and CagFbFP-Q148K (plum). Both structures show the same rotamer conformation of the lysine side chain, as well as a bending of the backbone. In addition, the respective buffer molecules of iLOV-Q489K and CagFbFP-Q148K are shown, namely the acetate ion (magenta) and the morpholine [O(CH_2_CH_2_)_2_NH](olive green). (D) Environment of the buffer molecules found in iLOV-Q48K and CagFbFP-Q148K, namely the acetate anion and in almost identical position the morpholine molecule. Both molecules form hydrogen bonds with the FMN, N468 (iLOV-Q489K) or N127 (CagFbFP-Q148K), respectively. Color coding as in panel C. Dashed lines represent hydrogen bonds with a donor-acceptor distance of ≤ 3.2 Å

While in parental iLOV the Iβ-strand is constituted by the residues 482 to 491, Iβ is shortened by two residues in iLOV-Q489K and ends at K489, with the remaining C-terminus kinking downwards relative to parental iLOV (Figure 2, A) with the C-terminal end of Iβ becoming detached from the core (in the following described as unlatching). In addition, the last three C-terminal residues lack detectable density. Interestingly, D491 and G492 have undergone post-translational modification to yield a cyclic imide structure (succinimide), which we attribute as an artifact due to protein aging [23]. Relative to Q489, that in parental iLOV forms an H-bond with the FMN-O4 atom, the side chain of the introduced K489 adopts a conformation that is extended away from the FMN chromophore, and forms an H-bond with the O2 atom of the succinimide ring (K489-NZ^…^SNN-O2, 3.1 Å) (Figure 2, B). Collectively, this results in opening up of the LOV core domain (Figure 2, A and B), which is expected to result in the influx of solvent molecules. In fact, additional electron density is present close to FMN-N5, which we interpret as an acetate ion (Figure 2, B, Figure S1, A), likely derived from the crystallization screen containing acetate buffer. Interestingly, in iLOV-Q489K, this ion occupies the former position of the Q489 side chain of parental iLOV. The here experimentally observed K489 conformation is distinct from the ones observed in previous MD simulations, suggesting that the previously described MD-identified K489_out_ conformation [19, 20] represents a local energy minimum, while the final, experimentally determined structure was energetically not accessible likely due to too short simulation times.

Irrespective of those discrepancies, the introduced lysine adopts a conformation that is facing away from the chromophore, which, in light of previous quantum mechanics molecular mechanics (QM/MM) calculations [19, 20] accounts for the blue-shifted spectral properties of the variant. Increased solvent influx, as observed in our structures, might hereby have an impact the magnitude of the blue shift, as the stabilization of solvent/buffer molecules that interact with the flavin (similarly to Q489 in parental iLOV) might affect the effective spectral properties.

To verify, that the structural changes, which we observed for iLOV-Q489K are indeed a consequence of the introduced Q489K mutation, we obtained the structure of the corresponding CagFbFP-Q148K variant. Altogether, we were able to obtain three types of CagFbFP-Q148K crystals, all using the precipitant solutions with pH of 6.5 (data and refinement statistics are summarized in Supplementary Table S1). In all of the obtained models the C-terminus of the protein is clearly unlatched, and the side chain of the introduced lysine is facing away from FMN as was found in iLOV-Q489K (Figure 2, C). Thus, the succinimide modification present in iLOV-Q489K does not influence the backbone shape, because CagFbFP-Q148K lacks this modification. Interestingly, the highest-resolution structure (PDB-ID 6YX4, determined at a resolution of 1.36 Å; C-α atom RMSD to iLOV-Q489K of 0.7675 Å for 102 aligned residues), revealed a six-membered ring in a chair conformation close to the FMN (Figure 2, D; Figure S1, B), which we attributed to derive from the precipitant solution containing 2-(N-morpholino)ethanesulfonic acid (MES). We thereby assume that the MES molecule was degraded during storage of the precipitant solution or during crystallization, and morpholine bound to the pocket. Similar degradation of a related buffer component 3-(N-morpholino)propanesulfonic acid (MOPS) and binding of morpholine to the protein was observed by Sooriyaarachchi *et al*. during crystallization of *Plasmodium falciparum* 1-deoxy-d-xylulose-5-phosphate Reductoisomerase [24] (PDB-ID 5JO0). Interestingly, the protein also crystallized using another precipitant solution for which we obtained crystals in two crystal forms. P2_1_2_1_2 crystals (PDB-ID: 6YX6) diffracted anisotropically, and we applied ellipsoidal truncation (see Supporting Materials and Methods for details) using the resolution cut-offs of 1.5 and 2.05 Å. Despite of having been crystallized under different conditions and in a different space group, all alternative CagFbFP-Q148K structures showed an extended outward facing K148 conformation as well as an unlatched C-terminus relative to parental iLOV (exemplarily shown for the highest resolution structure, PDB-ID: 6YX4; Figure 2, C, D). Therefore, we conclude that irrespective of the buffer and post-translational modification the C-terminus of the iLOV-Q489K and CagFbFP-Q148K variants is unlatched, the LOV core domain is opened up to allow solvent ingress and the introduced lysine occupies an extended conformation facing away from the FMN chromophore. We thus suggest these structural changes as the characteristic features for a blue-shifted absorption and emission spectrum, i.e. when compared to the parental FbFPs.

As shown by our above presented iLOV-Q489K and CagFbFP-Q148K structures, introduction of the single Q489K/Q148K substitution results in blue-shifted spectra, likely due to the displacement of the introduced lysine away from the FMN chromophore. In contrast, previous computational studies have suggested that retention of a charged lysine close to the chromophore should result in red-shifted spectral properties[19, 21]. We therefore reasoned that secondary mutations are needed to stabilize the introduced lysine to stay close to the FMN ring system. Based on our iLOV-Q489K structure we selected G487 and V392 as suitable positions as they are close to the introduced K489 (Figure S1, C), so that side-chain alterations at those sites could stabilize K489 to retain its position close to FMN. Following this rationale, we set out to mutate G487 and V392 in iLOV-Q489K. To sample a larger sequence space and to increase the chances for identifying a red-shifted FbFP variant, we generated two site-saturation libraries of iLOV-Q489K. Overall, 465 clones were screened per library. The fluorescence-emission maxima of a subset of promising variants identified by the screen are given in Table S2. While saturation mutagenesis at position G487 yielded no variants with red-shifted emission relative to parental iLOV, two variants with red-shifted emission maxima relative to iLOV-Q489K (Table S2, Figure S2) were identified. Both variants turned out to contain a serine substitution at position G487 (Table S2). Site-saturation at position V392, yielded more promising results as overall 16 variants with red-shifted spectral properties could be identified (Table S2). 14 out of 16 variants carried a threonine at position 392, while two variants contained either a cysteine or an alanine (Table S2). We selected the iLOV-V392T-Q489K variant, with the largest red shift relative to parental iLOV (variant B5-B12; Table S2, 8 nm red shifted fluorescence emission), for further studies, and overexpressed and purified the protein. Absorption and fluorescence emission spectra of the purified protein are shown in Figure 3, A. Compared to parental iLOV (Figure 3, A and B; grey dashed line), the double mutant iLOV-V392T-Q489K possesses a very similar absorption maximum at around 447 nm (Figure 3, A; red solid line, Table 1). The main fluorescence-emission maximum is red shifted by 6 nm to 502 nm (Figure 3, B; red solid line; Table 1). Compared to iLOV-Q489K (Figure 3, A, B; blue solid line), the double variant shows a red-shifted absorption maximum and a fluorescence-emission maximum shifted by 15 nm (Figure 3, A, B; red solid line; Table 1). To verify the transferability of the mutation, we introduced the corresponding I52T mutation into CagFbFP-Q148K. The resulting purified protein showed very similar spectral properties as iLOV-V392T-Q489K, i.e. basically no shift in the absorption maximum relative to parental CagFbFP (Figure 3, C; compare solid red line and grey dashed line; Table 1) as well as a 7 nm red-shifted fluorescence-emission maximum (Figure 3, D; compare grey dashed line and solid red line; Table 1). To rule out, that the I52T mutation alone is sufficient to generate the observed red-shift, we introduced the I52T mutation in parental CagFbFP (Figure 3, C, D; light red solid line; Table 1), which shows nearly identical spectral properties as parental CagFbFP (Figure 3, C, D; grey dashed line; Table 1). The latter is corroborated by the X-ray structure of CagFbFP-I52T (see Table S1 for data and refinement statistics), which is virtually identical to parental CagFbFP (C-α atom RMSD 0.5241 Å, 102 aligned residues). Most importantly, it does not show an unlatched C-terminus as observed for CagFbFP-Q148K (Figure S1, D).

**Figure 3:**
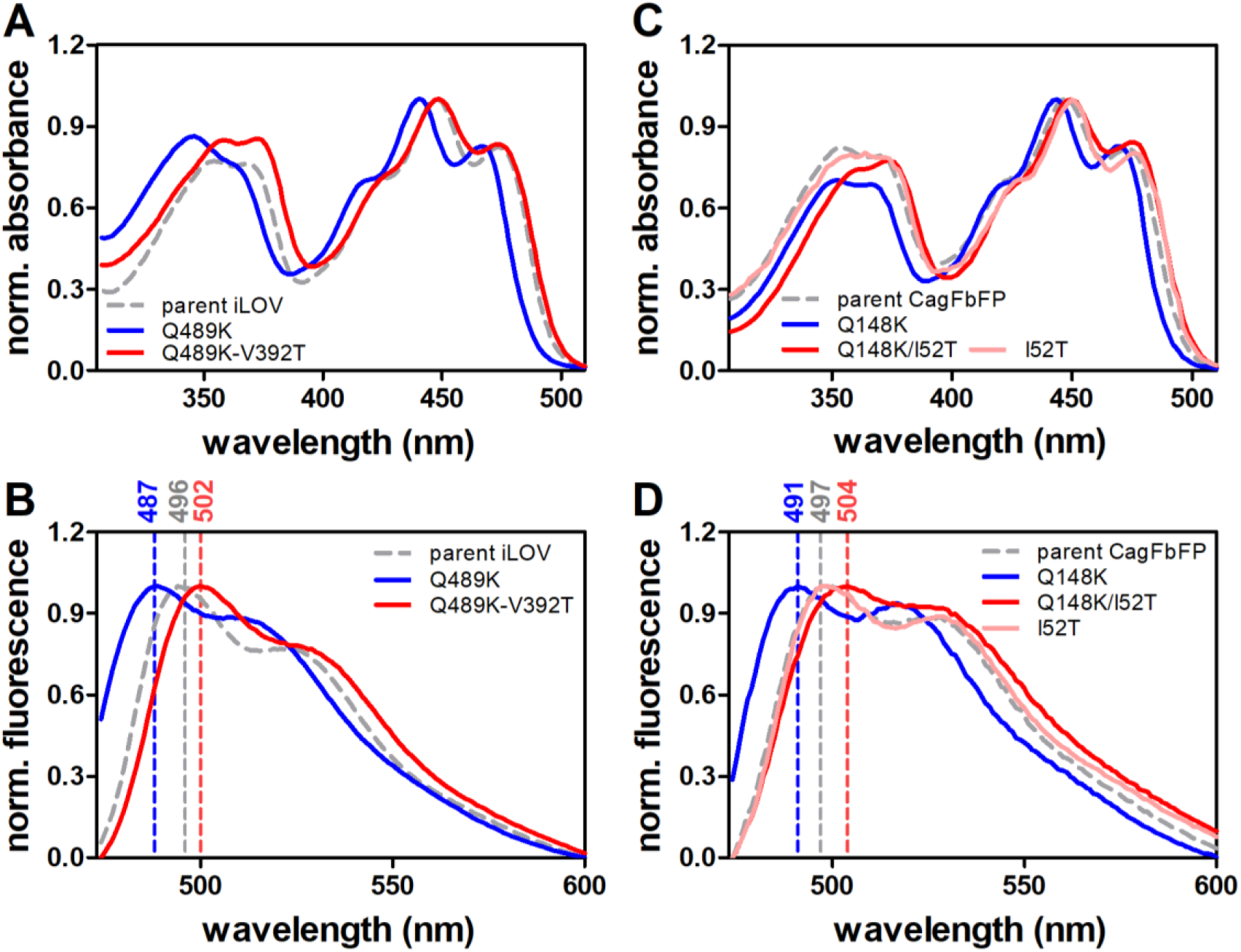
Absorption (A, C) and fluorescence-emission spectra (B, D) of iLOV and its variants (A, B) and CagFbFP and its variants (C, D). The spectra of the respective parental protein are shown as grey dashed lines, the blue-shifted variants (iLOV-Q489K; CagFbFP-Q148K) with blue solid lines and the red-shifted (iLOV-Q489K-V392T; CagFbFP-Q148K-I52T) with red solid lines. Additionally, the absorption and fluorescence-emission spectra of CagFbFP-I52T is shown (light red solid lines). Dashed lines of the respective color mark fluorescence emission maxima. All spectra are normalized to the corresponding maximum.

We thus conclude that a single secondary mutation in blue-shifted iLOV-Q489K and CagFbFP-Q148K, which places a polar threonine residue into the flavin-binding pocket, is sufficient to invert the spectral properties from blue-to red-shifted relative to the parent protein. To illustrate why until now the here identified FbFP variants were not identified by random mutagenesis or gene shuffling in various mutagenesis campaigns [1, 25, 26], we estimated the probability to identify the iLOV-V392T-Q489K from a large random error-prone PCR library. Here, when screening an error-prone PCR library having 10^6^ variants, generated with a medium mutation rate of 8 mutations/kb [27], the probability of discovering at least one copy of iLOV-V392T-Q148K variant is approximately 0.0014 (assuming roughly uniform mutation spectrum and independence across variants and DNA positions). Please note that for screening of 10^6^ clones already ultrahigh-throughput systems, such as FACS-sorting or droplet-based microfluidics screens [28] would be needed. Hence, ultrahigh-throughput methods with sufficient sensitivity to differentiate between blue- and red-shifted variants, as well as mutagenesis methods that e.g. allow the parallel saturation of multiple sites [29-31], or high-error rate ep-PCR methods [32], would likely be needed for further color-tuning of FbFPs.

To validate our design approach and elucidate the structural basis for the red-shifted spectral properties of the corresponding iLOV and CagFbFP variants, we tried to crystallize both variants. We readily obtained orthorhombic crystals for CagFbFP-I52T-Q148K (space group P2_1_2_1_2, two molecules per asymmetric units) that diffracted up to 1.8 Å. Compared to the blue-shifted iLOV-Q489K and CagFbFP-Q148K variants, the red-shifted double variant shows a closed FMN binding pocket with no unlatched C-terminus (Figure 4, A; ; C-α atom core RMSD: 0.4782 Å with 102 aligned residues).

**Figure 4:**
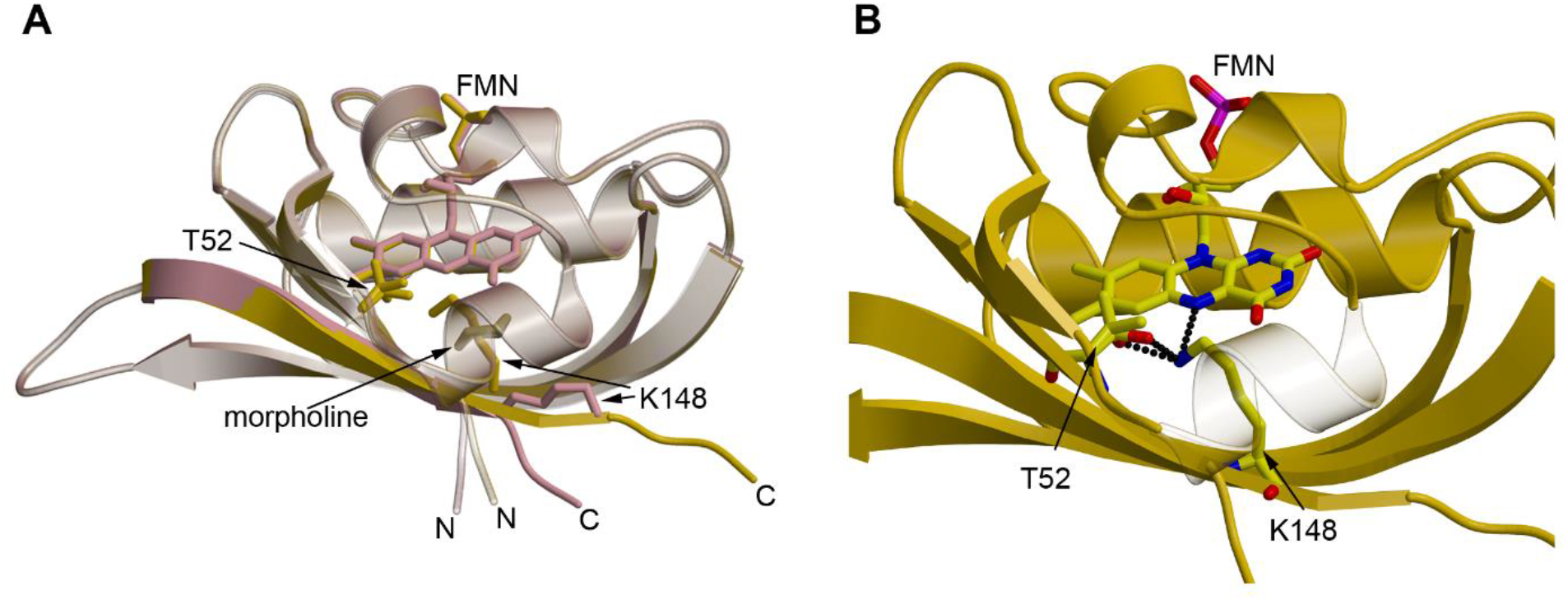
Structure of CagFbFP-I52T-Q148K. A) Superposition of CagFbFP-Q148K (plum) with CagFbFP-I52T-Q148K (gold). In addition, the introduced K148 and T52 (with two rotamer conformations) are depicted. B) Structure of CagFbFP-I52T-Q148K with hydrogen bonds between K148, T52 and the FMN. T52 shows two clear conformations, this is conditionally also to be observed for lysine, but for reasons of clarity only one conformation is shown. Additional details and the corresponding electron density maps are shown in Figure S3. Dashed lines represent hydrogen bonds with a donor-acceptor distance of ≤ 3.2 Å

The introduced T52 shows two conformations in both subunits, one with the side-chain oxygen facing towards G146 and a second one with the side-chain oxygen facing towards F69 (approximate occupancies of the two conformations are 65% and 35%, respectively). K489 adopts one major conformation in both subunits (approximate occupancy 65%), in which the side chain extends towards the introduced T52 and the FMN moiety. Interactions between both threonine conformations and the lysine are feasible with interatomic distances between the lysine NZ atom and the threonine side-chain oxygen of 2.5 or 2.6 Å in both subunits (Figure 4, B). It is remarkable that the double mutation (CagFbFP-I52T-Q148K) forces the lysine into a new conformation by hydrogen bonding with threonine, thus preventing the bending of the C-terminus, thereby restoring a nearly “native” closed LOV-core conformation.

In chain A, the introduced lysine adopts a second conformation with its side chain facing away from T52 pointing instead towards N127 and F125 (occupancy 35%) (Figure S3). This indicates that both of the introduced residues show a certain degree of flexibility. Stressing this notion, other residues such as D55, Q112, R119, D121, Q123, Q128, S130, S132, V145 show multiple conformations and two backbone conformations are observed in chain B for residues following V147. In conclusion, the here obtained CagFbFP-I52T-Q148K structure is in full agreement with our design concept, with the introduced threonine stabilizing the lysine in a conformation close to the FMN-N5 atom (interatomic distances 3.2 and 3.0 Å in chain A and B, respectively) (Figure 3, B).

To gain a better understanding of the photophysics of the different variants we determined their fluorescence quantum yields *Ф*_*F*_ and fluorescence lifetimes *τ*_*av*_ (Table 1). The corresponding fluorescence decays showed biexponential decay behaviour and are shown in Figure S4 and S5, respectively. The first unexpected finding are the almost identical photophysical parameters of the two parental proteins, CagFbFP and iLOV, respectively. Absorption and emission maxima, fluorescence quantum yields and fluorescence lifetimes are virtually identical (see Table 1). This is quite remarkable since origin and amino acid sequence of the two proteins are rather different and FbFPs *Ф*_*F*_ and *τ*_*av*_ are known to vary by more than 50% [18].

The photophysical properties of the CagFbFP variants are relatively straightforward to interpret. Compared to the parental CagFbFP protein (*Ф*_*F*_ = 0.35 ± 0.01; *τ*_*av*_ = 4.53 ± 0.03 ns) we observed a significant reduction of *Ф*_*F*_ along with similarly shorter *τ*_*av*_ for CagFbFP-Q148K (*Ф*_*F*_ = 0.24 ± 0.01; *τ*_*av*_ = 3.28 ± 0.03 ns), which can be attributed to dynamic quenching processes by solvent molecules, i.e. due to increased solvent access, as revealed by our structural analyses (see above). This scenario is corroborated by the data obtained for CagFbFP-I52T, which is structurally identical to the parent protein and showed, within the experimental error very similar *Ф*_*F*_ and *τ*_*av*_ values (*Ф*_*F*_ = 0.38 ± 0.06; *τ*_*av*_ = 4.42 ± 0.07 ns). The same holds for, CagFbFP-I52T-Q148K, which possesses a closed FMN-binding pocket with K148 being in H-bonding distance to the FMN-N5 atom, and consequently shows increased *Ф*_*F*_ and *τ*_*av*_, (*Ф*_*F*_ = 0.27 ± 0.01; *τ*_*av*_ = 3.82 ± 0.03 ns) relative to CagFbFP-Q148K. Please note however, that CagFbFP-I52T-Q148K does not show a completely parental-CagFbFP-like *τ*_*av*_ value, hinting at a certain degree of flexibility or conformational heterogeneity in this variant, which is indeed observed in the CagFbFP-I52T-Q148K structure (multiple conformations for I52 and K148 as well as multiple backbone conformations for the C-terminus in chain A; see above).

Surprisingly, even though having very similar spectral properties and overall structures, the situation is less clear for iLOV and its variants. Compared to parental iLOV (*Ф*_*F*_ = 0.33 ± 0.01), iLOV-Q489K possesses a very similar *Ф*_*F*_ (*Ф*_*F*_ = 0.35 ± 0.01), while *τ*_*av*_ values for iLOV-Q489K are shorter (*τ*_*av*_ = 4.03 ± 0.08 ns) as compared to the parental protein (*τ*_*av*_ = 4.47 ± 0.08 ns). This trend cannot be explained by dynamic quenching processes, which occur in the excited state and hence influence both *Ф*_*F*_ and *τ*_*av*_ in a similar fashion. To complicate matters, similar to the situation for the CagFbFP-I52T-Q148K variant, the corresponding iLOV-V392T-Q148K variant possesses virtually identical *Ф*_*F*_ and *τ*_*av*_ (*Ф*_*F*_ = 0.33 ± 0.02; *τ*_*av*_ = 4.48 ± 0.03 ns) as the parent protein. Thus, either the iLOV-Q489K variant or the parent iLOV protein shows divergent behavior as compared to the parent CagFbFP protein. Possible explanations include a change in the electronic structure of the first excited singlet state (perhaps also the ground state) of iLOV-Q489K, which manifests in blue-shifted absorption and emission spectra and an in total faster deactivation of the first excited singlet state with larger relative contribution from fluorescence. In addition, altered photochemical properties e.g. increased light-dependent formation of the neutral FMN semiquinone radical FMNH· [33] in the parent protein or – more simple – an increase in rate of internal conversion relative to iLOV-Q489K and concomitantly a decrease in *Ф*_*F*_, could also account for the discrepancy between the iLOV variants and their CagFbFP counterparts.

To computationally probe the hypothesis that the experimentally observed lysine conformations of the blue- and red-shifted variants are the cause for the observed spectral properties, we performed QM/MM calculations based on the corresponding crystal structures. From the calculated vertical excitation energies, we observed that the calculated absorption and emission wavelengths follow the same general trend, i.e. with regard to the observed direction of the spectral shift, as the experimental ones (see Table S3). Our calculations thus support the idea that the Q489K/Q148K substitutions, and hence the resulting structural changes (Iβ unlatching) in iLOV and CagFbFP are the cause for blue-shifted absorption and fluorescence emission with respect to the parental proteins. The double substitutions (Q489K/Q148K and V392T/I52T) in the two proteins in turn lead to red-shifted absorption and fluorescence emission, by stabilizing the introduced lysine in a position close to the FMN chromophore.

In many physiology and cell biology studies it is necessary to observe simultaneously many proteins and cell parameters, where the number of spectrally different detection channels is often a limiting factor. In commercial fluorescence microscopes two, three or four spectrally resolved channels are available. The new FbFP variants show similar but significantly different fluorescence spectra and lifetime differences, which can be exploited in fluorescence microscopy applications utilizing the same excitation wavelength and emission detection channel. A separation of two FbFPs by spectrum or lifetime difference excited with the same laser and observed in the same detection channel adds an additional detection channel, e.g. one more protein can be observed. We wanted to demonstrate the usefulness of the new FbFP mutants for this purpose using two of them and chose a blue and red-shifted FbFP mutant as example. To this end, we transformed the *E. coli* cells with plasmids carrying the genes for CagFbFP-Q148K (blue-shifted variant) and CagFbFP-I52T-Q148K (red-shifted variant). Compared to untransformed cells, both new strains revealed bright fluorescence (Figure 5, A-D). When mixed, the cells of the two strains - though not unequivocally distinguishable in the fluorescence intensity image (even with the spectral dimension used in the image presentation) - can clearly be differentiated from each other using spectral unmixing (Figure 5, E,F). Therefore, the identified mutants allow for two-color microscopy with a single excitation wavelength and observed in the same spectral range. Instead of distinguishing different cell organelles or different proteins in mammalian cells.

**Figure 5:**
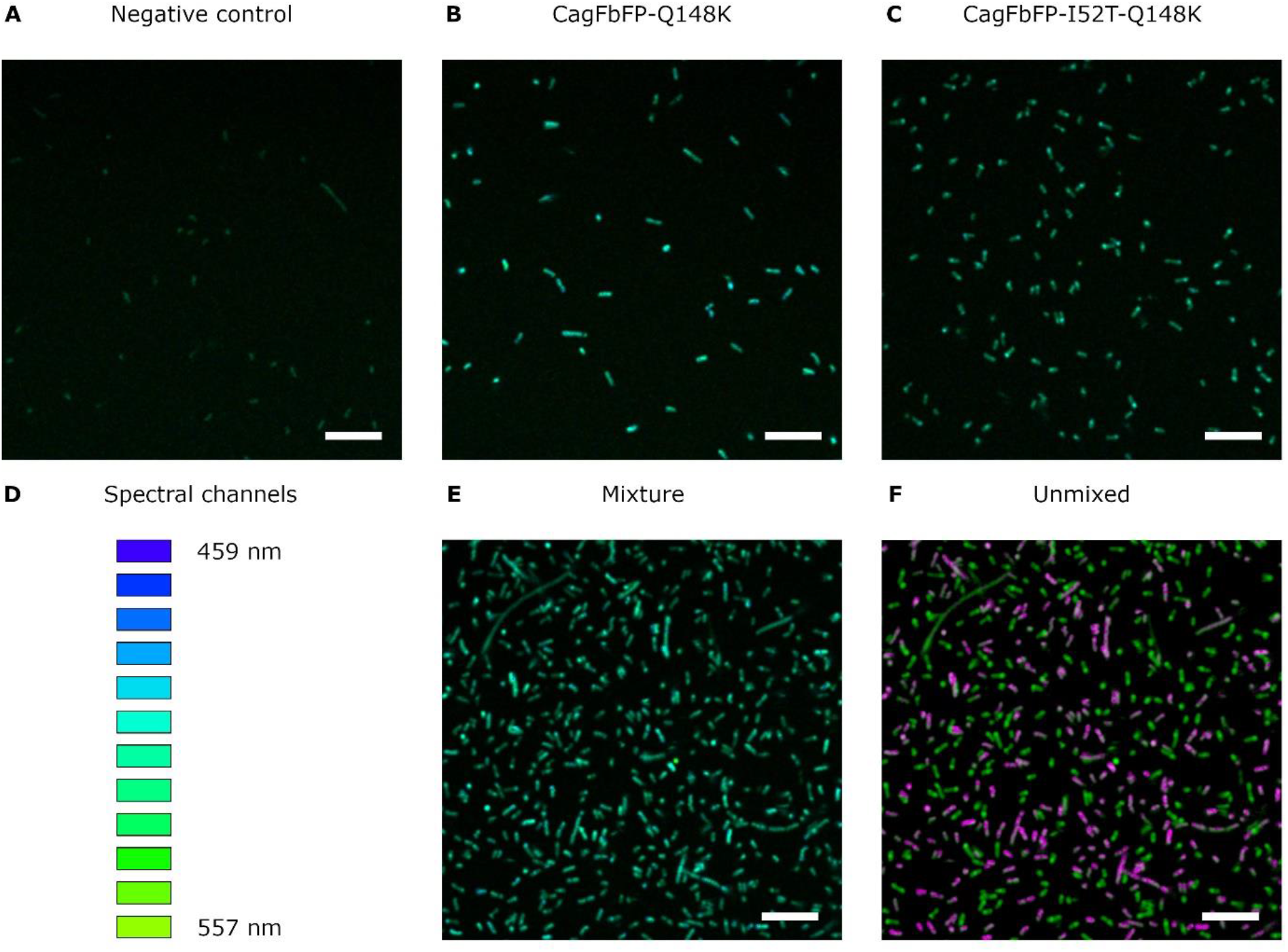
Imaging of *E. coli* cells transformed with different FbFP variants. Combined fluorescence intensity and fluorescence spectrum representation of the emission in the pixels is shown. A) Untransformed *E. coli* cells used as negative control. B) *E. coli* cells expressing CagFbFP-Q148K. C) *E. coli* cells expressing CagFbFP-I52T-Q148K. D) Diagram showing the perceived colors for different fluorescence acquisition channels. Twelve channels from 459 to 557 nm with 9 nm step were used. E) Mixture of *E. coli* cells expressing CagFbFP-Q148K and *E. coli* cells expressing CagFbFP-I52T-Q148K. F) Same as E) processed using linear unmixing. The cells expressing CagFbFP-Q148K are highlighted in magenta and the cells expressing CagFbFP-I52T-Q148K are highlighted in green. All images except for F) were obtained using the same hardware and software settings. The scale bar is 10 µm

## Conclusions

The presented work provides a firm structural understanding of spectral tuning in flavin-based fluorescent proteins (FbFPs), an emerging class of fluorescent reporters. Simultaneously, the presented data improves our understanding of the photophysics and the tunability of the spectral properties for flavins in general, which are essential cofactors in a variety of enzymatic systems and various photoreceptor families. Finally, the generated blue- and red-shifted variants represent promising fluorescent reporters with photophysical properties similar to their respective parent proteins and hence should find application as oxygen-independent reporters in life sciences. Although, the blue- and red-shifted variants are spectrally not very different, differentiation between the two proteins is possible in combination with modern fluorescence techniques like spectral unmixing, a commercially available technique with high-end confocal fluorescence microscopes. This combination opens up the possibility of multicolor imaging in anaerobic settings, where GFP and related proteins do not work due to their oxygen-dependent fluorophore maturation [2]. In addition, the large fluorescence lifetime difference between CagFbFP and its Q148K variant allows similar differentiation of two bacteria or proteins by fluorescence lifetime imaging with only one excitation wavelength.

### Experimental procedures

#### Cloning, site directed and site-saturation mutagenesis of iLOV

Site saturation at position 392 and 487 of the iLOV encoding gene (*Arabidiopsis thaliana*, Phot1 numbering) was achieved by two single-primer reactions performed in parallel, which were, after initial amplification, combined to allow site saturation using a Quikchange PCR approach [34]. Oligonucleotide primers were designed using a degenerate NNS codon instead of the parental target codon. The initial two parallel reactions contained pET28a-iLOV-Q489K [19] as a template with 0.5 µM iLOV-V392X_fw: 5’-GC CAT ATG ATA GAG AAG AAT TTC NNS ATC ACT GAT CCT AGG CTT CCC GA −3’) and iLOV-V392X_rev: 5’-T CGG GAA GCC TAG GAT CAGT GAT SNN GAA ATT CTT CTC TAT CAT ATG GC −3’) or the corresponding iLOV-G487X_fw: 5’-G GGA GAG CTT CAA TAC TTC ATC NNS GTG AAA CTC GAT GGA AGT GAT C −3’ and iLOV-G487X_rev: 5’-G ATC ACT TCC ATC GAG TTT CAC SNN GAT GAA GTA TTG AAG CTC TCC C −3’ primer, respectively, 0.2 mM dNTPs, 1x GC-buffer and Phusion^®^ High-Fidelity DNA polymerase (Thermo Fisher Scienitfic, Waltham, MA USA). Amplification was achieved using the following protocol: initial denaturation for 3 min at 98 °C, followed by 5 cycles of denaturation at 98 °C for 3 min, annealing at 64 °C for 0.5 min, elongation at 72 °C for 6 min. After five cycles the two separate reactions were combined and additional 20 cycles were carried out using the same temperature program followed by a final elongation step at 72 °C for 10 min. Methylated parental template DNA was digested with *Dpn*I. The reaction was stopped at 75 °C for 20 min. The resulting plasmid saturation libraries were designated as pET28a-iLOV-Q489K-V392X and pET28a-iLOV-Q489K-AG487X.

#### Cloning and site-directed mutagenesis of CagFbFP

The mutation Q148K was introduced into the parental CagFbFP [22] encoding gene using two PCR reactions. In the first reaction, the gene fragment corresponding to amino acids 47-147 of the parent protein was amplified using pET11-CagFbFP as a template with 0.5 μM CagFbFP-fw: 5’-TAT ACA TAT GGC CAG CGG TAT GAT TGT T-3’ and CagFbFP-147-rev: 5’-CAC CAA CAA ATG CAA CAA CAT TGC C −3’ primers, 0.2 mM dNTPs, 1x KAPA2G Buffer A and KAPA2G Robust DNA Polymerase (Merck KGaA, Darmstadt, Germany). Amplification was achieved using the following protocol: initial denaturation for 3 min at 95 °C, followed by 30 cycles of denaturation at 95 °C for 15 sec, annealing at 60 °C for 15 sec and elongation at 72 °C for 15 sec. Final elongation step at 72 °C was for 10 min. This gene fragment was excised from the gel, purified using the Cleanup Standard kit (Evrogen, Mosow, Russia). A second extension PCR reaction was performed using the purified fragment, an additional 0.02 μM oligonucleotide with mutation CagFbFP-Q148K-fw: 5’-GGC AAT GTT GTT GCA TTT GTT GGT GTT AAA ACA GAT GTT ACC GCA CAT CAT CAT C −3’ and 0.5 μM and primers CagFbFP-fw: 5’-TAT ACA TATG GCCA GCG GTA TGA TTG TT-3’ and CagFbFP-rev: 5’-AGC CGG ATC CTT AGT GAT GGT GAT G −3’, 0.2 mM dNTPs, 1x KAPA2G Buffer A and KAPA2G Robust DNA Polymerase (Merck KGaA, Darmstadt, Germany). Amplification was achieved using the same protocol. The resulting gene with the Q148K mutation was excised from the gel, purified using the Cleanup Standard kit (Evrogen, Moscow, Russia). The restriction reaction was carried out using *BamH*I and *Nde*I restriction enzymes. The resulting sticky ends were ligated to the vector pET11 using T4 DNA Ligase (Thermo Fisher Scienitfic, Waltham, MA USA). The mutation I52T was introduced into CagFbFP and CagFbFP-Q148K using PCR. The reaction mixture contained pET11-CagFbFP or pET11-CagFbFP-Q148K as a template with 0.5 µM CagFbFP-I52T_fw: 5’-CGG TAT GAC CGT TAC CGA TGC CGG - 3’ and CagFbFP-I52T_rev: 5’-CGGTAACGGTCATACCGCTGGCC - 3’ primers, 0.2 mM dNTPs, 1x Phusion HF buffer and Phusion® High-Fidelity DNA polymerase (Thermo Fisher Scienitfic, Waltham, MA USA). Amplification was achieved using the following protocol: initial denaturation for 1 min at 98 °C, followed by 25 cycles of denaturation at 98 °C for 10 sec, annealing at 55 °C for 0.5 min and elongation at 72 °C for 2.5 min. Final elongation step at 72 °C was for 10 min. Methylated parental template DNA was digested with *Dpn*I. The reaction was stopped at 80 °C for 5 min. Plasmids pET11-CagFbFP-I52T and pET11-CagFbFP-Q148K-I52T were transformed into *E. coli* DH10B cells, plated on LB agar supplemented with ampicillin (100 µg /ml) and incubated over night at 37 °C. The plasmid DNA of the CagFbFP variants was isolated using the Plasmid Miniprep kit (Evrogen, Moscow, Russia). The resulting mutated CagFbFP encoding DNA insert was subsequently sequenced (Evrogen, Moscow, Russia) to verify the introduced amino acid exchange.

### Expression and high-throughput screening of the iLOV-Q489K-V392X site saturation library

*E. coli* BL21(DE3) cells were transformed with the pET28a-iLOV-Q489K-V392X and pET28a-iLOV-Q489K-G487X libraries, plated on LB agar supplemented with kanamycin (50 µg/µL), and incubated over night at 37 °C. To obtain a pre-culture, each well of a 96-deep well plate, filled with 500 µL LB supplemented with kanamycin, was inoculated with a single colony. Three wells of each plate were used as controls, containing either non-inoculated medium, *E. coli* BL21(DE3) pET28a-iLOV or pET28a-iLOV-Q489K. The deep well plates were then incubated at 37 °C and 800 rpm for 16 h. Subsequently, 500 µL of modified auto-induction media (AIM) [18, 35] were inoculated with the pre-culture using a microplate replicator and incubated at 37 °C and 800 rpm for 16 h. Cells were harvested by centrifugation at 2000 rpm for 30 min and lysed using BugBuster® protein extraction reagent (Merck, Darmstadt, Germany) following the instructions provided by the manufacturer. After centrifugation at 2000 rpm, 100 µL of the cell free, protein containing supernatant were transferred to a black 96 well microtiter plate (NUNC, Thermo Fischer Scientific, Darmstadt, Deutschland) and emission spectra were recorded using an Infinite 1000 PRO microplate reader (Tecan, Crailsheim, Germany; excitation wavelength, λ_ex_ = 440 nm; emission wavelength range, λ_em_ = 456 − 600 nm; slit width of 10 nm). Overall, five plates a’ 93 clones (+ 3 controls), resulting in 465 individual clones per library, were screened. The plasmid DNA of the iLOV variants, identified in the screen to possess spectrally red-shifted fluorescence emission, was isolated from replica plates using the innuPREP Plasmid Mini Kit 2.0 (Analytik Jena, Germany). The resulting mutated iLOV encoding DNA insert was subsequently sequenced (Seqlab, Göttingen, Germany) to identify the introduced amino acid exchange.

### Expression and purification of iLOV and its variants

Verified plasmids (pET28a-iLOV, pET28a-iLOV-Q489K, pET28a-iLOV-Q489K-V392T) were transformed into *E. coli* CmpX131 [36]. The corresponding proteins were produced and purified by immobilized metal-ion affinity chromatography (IMAC), as described previously [19]. After IMAC purification the imidazole-containing elution buffer was exchanged for storage buffer (10 mM Tris/HCl, pH = 7.0, supplemented with 10 mM NaCl) by using a G25 Sephadex desalting column. Protein samples were concentrated using Macrosep® advance centrifugal device (PALL, Dreieich, Germany; molecular weight cut-off = 3 kDa) and stored at 4 °C in the dark.

### Expression and purification of CagFbFP and its variants

Verified plasmids (pET11-CagFbFP, pET11-CagFbFP-Q148K, pET11-CagFbFP-I52T, pET11-CagFbFP-Q148K-I52T) were transformed into *E. coli* C41(DE3) or *E. coli* BL21(DE3). The corresponding proteins were produced and purified by immobilized metal-ion affinity chromatography (IMAC), as described previously [22]. After IMAC purification the imidazole-containing elution buffer was exchanged for storage buffer (10 mM sodium phosphate, pH = 8.0, supplemented with 10 mM NaCl) by using size-exclusion chromatography on a Superdex® 200 Increase 10/300 GL column (GE Healthcare Life Sciences, USA) or by using PD10 desalting columns (GE Healthcare Life Sciences, USA). Protein samples were concentrated using Amicon® Ultra-4 Centrifugal Filter Unit (Merck KGaA, Darmstadt, Germany; molecular weight cut-off = 10 kDa) and stored at 4 °C in the dark.

### Fluorescence spectroscopy

Fluorescence emission and excitation spectra were recorded using a Quanta-Master 40 fluorescence spectrophotometer (Photon Technology International, Birmingham, NJ, USA) as described previously by Wingen and co-workers [18]. Absorbance spectra were recorded using Cary 60 UV/Vis spectrophotometer (Agilent Technologies, Santa Clara, CA, USA) equipped with a Peltier-thermostated single cell-holder. Fluorescence quantum yields were determined using a Quanta-Master 40 fluorescence spectrophotometer (Photon Technology International, Birmingham, NJ, USA) equipped with an integrating sphere as described in Wingen *et al*. [18]. All measurements were carried out in storage buffer (10 mM Tris/HCl, pH = 7.0, supplemented with 10 mM NaCl; iLOV and its variants, or in 10 mM sodium phosphate, pH = 8.0, supplemented with 10 mM NaCl; CagFbFP and its variants).

### Fluorescence lifetime measurements

The time-resolved detection of the fluorescence intensity decay of all FbFPs was performed using a Fluotime100 fluorescence spectrophotometer (Picoquant, Berlin, Germany) based on a picoHarp300 unit by using a pulsed diode laser (Laser Picoquant LDH-C440; emission: 440 nm; pulse width: 50 ps; used repetition frequency: 20 MHz) as an excitation source. Fluorescence decay curves as a function of time (t) were measured by time-correlated single-photon counting that enables the determination of fluorescence decay components with fluorescence lifetimes greater than 100 ps [37, 38]. Decay curves were analyzed by iterative reconvolution of the instrument response function, IRF(t), with an exponential model function, M(t), using the FluoFit software (version 4.5.3.0; Picoquant) applying eqn (1) and (2):

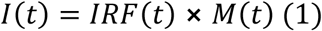

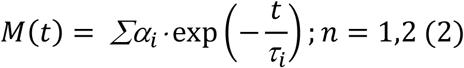

τ_i_ is the characteristic lifetime and α_i_ is the respective intensity. The average fluorescence lifetime, τ_fl,ave_, was calculated using eqn (3):

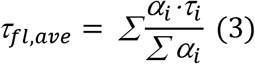

### Protein crystallization

The purified iLOV-Q489K protein (in 10 mM Tris/HCl, pH = 7.0, supplemented with 10 mM NaCl) was concentrated to 26 mg/mL using VivaSpin centrifugal concentrators (Sartorius, Göttingen, Germany; 20 kDa molecular-weight cut-off). Crystallization setups were performed using the sitting drop vapor diffusion method (1 μL of protein and 1 μL of the reservoir solution) and stored at 19 °C in the dark. Tetragonal crystals grew within 6 weeks against 0.1 M sodium acetate (pH 4.6) containing 25 % (w/v) PEG 1000. The purified CagFbFP variants (in 10 mM sodium phosphate, pH = 8.0, supplemented with 10 mM NaCl) were concentrated to 30 mg/ml using Amicon® Ultra-4 Centrifugal Filter Unit (Merck KGaA, Darmstadt, Germany; molecular weight cut-off = 10 kDa). Crystallization setups were performed using the sitting drop vapor diffusion method (150 nL of protein and 100 nL of the reservoir solution) and stored at 22 °C in the dark. CagFbFP-Q148K crystals were obtained using the precipitant solution containing 0.1 M MES monohydrate pH 6.5 and 12% w/v PEG 20000. Morpholine-free CagFbFP-Q148K structure was obtained using 0.5 M ammonium sulfate, 1 M lithium sulfate monohydrate, 0.1 M sodium citrate tribasic dihydrate pH 5.6. CagFbFP-I52T crystals were obtained using the precipitant solution containing 0.2 M lithium sulfate, 0.1 M BIS-Tris pH 5.5 and 25% PEG 3350. CagFbFP-I52T-Q148K crystals were obtained using the precipitant solution containing 0.1 M citrate pH 5.0 and 20% PEG 6000.

### Data collection and structure determination

iLOV-Q489K crystals grew in high PEG concentrations and were cryo-cooled directly at 100 K. X-ray diffraction data was recorded at the beamline ID29 (ESRF, Grenoble, France [39]) tuned to a wavelength of 0.91 Å on a PILATUS 6M-F detector (Dectris Ltd., Baden, Switzerland). Data collection strategy was based on calculations using the program BEST which accounts for radiation damage and symmetry [40]. Data processing was carried out using the program XDS [41] and AIMLESS (part of the CCP 4 package [42]). The initial phases were obtained by molecular replacement using MOLREP [42], and the search model was generated from the crystal structure of iLOV (PDB-ID: 4EES). Analysis of Matthews coefficient [42] suggested one molecule per asymmetric unit, with Matthews coefficient of 1.82 Å^3^/Da and the solvent content of 32.6 %. The model was further improved with several cycles of refinement using the PHENIX package [43] and manual rebuilding using the program COOT [44]. Data collection and refinement statistics are listed in Table S1.

CagFbFP crystals were cryoprotected by adding the precipitant solution (see above) supplemented with 20% glycerol and were cryo-cooled at 100 K. X-ray diffraction data for the different CagFbFP crystals was recorded at various beamlines (see Table S1 for details). Data collection strategy was based on calculations using the program BEST which accounts for radiation damage and symmetry [40]. Data processing was carried out using the program XDS [41] and AIMLESS (part of the CCP 4 package [42]). The dataset 6YX6 (see Table S1) revealed strong anisotropy and was processed using STARANISO [45] using the following criterion for determining the diffraction-limit surface: <Imean/σImean> of more than 2. The initial phases were obtained by molecular replacement using MOLREP [42], and the search model was generated from the crystal structure of CagFbFP (PDB-ID: 6RHF). The model was manually refined using the programs COOT [44] and REFMAC5 [46]. Data collection and refinement statistics are listed in Table S1.

### QM/MM calculations

QM/MM optimizations and spectral calculations were performed with the program package ChemShell v3.5 [47]. We used Turbomole 6.3 [48] for the QM part and DL_POLY [49] as driver of the Amber [50] force field for the MM part. Prior to QM/MM calculations force-field based energy minimization were carried out using Amber14 program [51] with the Amber ff99SB [52, 53] all-atom force field for proteins and general Amber force field (GAFF) [50] for flavin mononucleotide (FMN) and the TIP3P model for water [54]. We used the available atomic charges and force-field parameters for FMN reported in our previous work [19]. The initial coordinates for simulations were taken from X-ray structure of iLOV (PDB-ID: 4EES; [25]), iLOV-Q489K (PDB-ID: 7ABY), and parental CagFbFP (PDB-ID: 6RHF; [22]), CagFbFP-I52T-Q148K (PDB-ID: 7AB7). iLOV proteins contain one iLOV molecule per asymmetric unit and thus represents a monomeric structure, while CagFbFP variants have two CagLOV molecules per asymmetric forming a homodimeric structure. The initial coordinates of chain A of CagFbFP-Q148K-I52T structure were used for all simulations. The protonation states of titratable residues at pH 7 were assigned on the basis of pKa calculations using the PROPKA 3.1 program [55] and visual inspection. In simulations all charged residues were kept at their standard protonation states. Side chains of Asn and Gln residues were checked for possible flipping. To neutralize the systems, solvent water molecules that were at least 5.5 Å away from any protein atoms were replaced by Na^+^ ions. Hydrogen atoms were added employing the tleap module of AmberTools14 [51]. The protein was solvated in a water box of 12 Å radius centered at the center of mass. Crystal water molecules were kept. To calculate the electrostatic interactions a cutoff of 10 Å was used. The total size of the simulation systems were ∼ 27,000 atoms, including ∼8,500 TIP3P [54] water molecules. The solvent and the ions followed by the whole system were subjected to minimization using 10,000 steps of steepest descent followed by 3,000 steps of conjugate-gradient minimization. The minimized structures from the energy minimization runs were chosen as starting structures for further QM/MM calculations. For the QM part of the QM/MM calculation, time-dependent density functional (TD) with B3LYP density functional [56-58] and the SVP basis set [56-58] from the Turbomole basis set library has been chosen to obtain vertical excitation and emission energies. Reliable excitation energies have already been obtained by this combination of method and basis set for electronic states of the FMN in LOV domain [19]. Geometries of the ground state (S_0_) and first singlet excited state (S_1_) were optimized at QM/MM DFT/AMBER level of theory. We have applied the additive QM/MM scheme with electrostatic embedding. The charge-shift scheme [59] and hydrogen link atoms were employed to handle the QM-MM boundary region. During the QM/MM structure optimizations which were carried out using the DL-FIND [60] optimizer module of ChemShell, we relaxed all fragments (amino acids, FMN, water, counter ions) that lie within 10 Å of the FMN chromophore. Everything beyond this selection was kept frozen during QM/MM structure optimizations. The QM region consists of the FMN chromophore while the covalent bond (between C13 and C15 for FMN) across the QM-MM boundary was capped with hydrogen link atoms. VMD [61], and AmberTools 14 [51] were used for molecular visualizations and analysis of simulations.

### Microscopy

*E. coli* C41(DE3) cells were grown in a shaker at 37 °C. Expression was induced using 1 mM IPTG when the optical density of 0.6 was reached. Expression of CagFbFP-Q148K and CagFbFP-Q148K-I52T was continued for 100 minutes and 5 hours, respectively. The cells were harvested by centrifugation at 4000 g for 3 min. The pellets were resuspended in the phosphate-buffered saline buffer and diluted to the optical density of approximately 0.5 and placed on cover slips for imaging. The lambda stacks (459-557 nm, 9 nm step) were obtained using the LSM780 system (Carl Zeiss, Jena, Germany) with 100x objective (oil-immersive, NA=1.46). Fluorescence was excited by pulsed 900 nm Insight DeepSee laser (20 mW at the objective). Images were obtained with the resolution of 2048×2048 pixels, pixel dwell time of 3.1 µs and 4x line averaging. Afterwards, the images were averaged using 5 pixel moving average to further reduce spectral variation between neighboring pixels (caused by noise). The linear unmixing was performed with the ZEN software (Carl Zeiss, Jena, Germany) utilizing the fluorescence emission spectra obtained from images of cells expressing only either CaFbFP-Q148K or CagFbFP-Q148K-I52T.

## Data availability

All data described in the manuscript is contained within the main manuscript and supporting information. Protein crystal structures have been deposited under the following accession numbers (see also Table S1 for details): iLOV-Q489K: PDB-ID: 7ABY; CagFbFP-Q148K: PDB-ID: 6YX4, 6YX6, 6YXB; CagFbFP-I52T: PDB-ID: 7AB6; CagFbFP-I52T-Q148K: 7AB7

## Acknowledgements

We thank Yuval Nov (Department of Statistics, University of Haifa, Israel) for his help in estimating the probability of finding the ilOV-V392T-Q489K variant by error-prone PCR and Anastasia Smolentseva for help with expression and purification of CagFbFP variants.

## Funding and additional information

M.D.D thanks DFG Training Group 1166 “BioNoCo” (Biocatalysis in Non-conventional Media”) for the financial support and computing resources granted by JARA-HPC from RWTH Aachen University under project JARA0065. Part of this work was funded by the Federal Ministry of Education and Research (BMBF) in the framework of the collaborative research project “OptoSys” (FKZ 031A16 and FKZ 031A167B). Studies of CagFbFP-I52T were funded by RFBR, project number 20-34-70109. We acknowledge the Structural Biology Group of the European Synchrotron Radiation Facility (ESRF, Grenoble, France) for granting access and providing assistance in using the beamline ID29, and the Paul Scherrer Institut (PSI, Villigen, Switzerland) for provision of synchrotron radiation beamtime at beamline X06SA of the Swiss Light Source (SLS) and would like to thank Dr. Anuschka Pauluhn for assistance. We also acknowledge the European Molecular Biology Laboratory (EMBL) unit in Hamburg at Deutsche Elektronen-Synchrotron (DESY, Hamburg, Germany) for granting access to the synchrotron beamlines and are thankful to Dr. Gleb Bourenkov for technical assistance.

